# Metabolic Flexibility and Energy Substrate Utilization Regulate Contractility in the Human Myometrium

**DOI:** 10.64898/2026.02.02.702681

**Authors:** Kevin K. Prifti, Ritu M. Dave, Kaci T. Mitchum, Jordan L. Rich, Ruth M. Gill, Magdaleena N. Mbadhi, Antonina I. Frolova

## Abstract

The uterus requires energy for sustained contractility during labor, to deliver the fetus and diminish the risk of postpartum hemorrhage. Our objective was to define energy requirements and assess metabolic flexibility in quiescent and contractile myometrial cells. Cells were treated with oxytocin to stimulate myometrial contractility. We found that myometrial cells rely on oxidative phosphorylation during quiescence and, when treated with oxytocin, can adapt to higher energy demands by shifting their energy production to glycolysis. Treatment with mitochondrial oxidation inhibitors revealed that in quiescent myometrial cells basal oxygen consumption rate decreased when treated with glucose oxidation inhibitor UK5099, but not the long chain fatty acid oxidation inhibitor etomoxir or the glutamine oxidation inhibitor BPTES. In oxytocin treated myometrial cells, this decrease was also observed upon BPTES treatment in addition to UK5099, suggesting that contractile myometrial cells can shift energy production from glucose to glutamine. Functionally, myometrial contractility was significantly reduced by UK5099 but not by etomoxir, further indicating dependence on glucose utilization.

## INTRODUCTION

In labor, the uterus must contract with enough force, frequency, and endurance to deliver the fetus and diminish the risk of labor dystocia and postpartum hemorrhage. Like all cells, uterine smooth muscle (myometrial) cells rely on ATP synthesis for energy production, either through glycolysis or oxidative phosphorylation^1^. Although current understanding of myometrial energy requirements is limited, a few lines of evidence suggest that glucose is the primary energy source at term gestation. First, myometrium from pregnant, non-laboring women contains higher concentrations of glucose, a higher ratio of lactate to pyruvate, and lower triglyceride metabolite concentrations than skeletal muscle^2,3^. Second, studies comparing blood concentrations of energy substrates in uterine arteries vs. veins at term demonstrated a net uptake of glucose, but not glycerol or free fatty acids^4^. Third, in the rat uterus, glycogen stores increase by two-fold from early pregnancy to term^5^, suggesting that carbohydrates are stored in the uterus as it primes for delivery. In skeletal and cardiac muscles, metabolic flexibility, defined as the ability to shift between substrates or pathways in response to availability, is essential for maintaining function under altered metabolic demands or stress^6–8^. Whether the myometrium can adapt similarly and utilize alternative substrates to meet the high and sustained energy demands of labor is unknown.

In this study, our objective was to define energy requirements and assess metabolic flexibility in human myometrial cells *in vitro*. To model a contractile state, we treated cells with the nonapeptide hormone oxytocin. *In vivo*, oxytocin is released by the pituitary gland, binds to its receptor on myometrial cells, and triggers intracellular calcium influx^9–11^. This intracellular calcium flux activates calcium-calmodulin coupling, myosin light chain kinase, and ultimately the actin-myosin crossbridge that generates contractions^12^. In clinical practice, nearly half of all pregnant women in the United States are treated with oxytocin to induce or augment labor or to prevent postpartum hemorrhage^13,14^. We hypothesized that myometrial cells would alter their energy substrate preference following treatment with oxytocin. To assess metabolic activity, we used the Agilent Seahorse XF Analyzer to measure oxygen consumption rate as an indicator of oxidative phosphorylation and extracellular acidification rate as an indicator of glycolysis. To determine biologic relevance, we quantitate human myometrial tissue contractility *ex vivo* in presence of mitochondrial oxidation inhibitors. We present evidence that myometrial cells primarily rely on oxidative phosphorylation during quiescence and shift their energy production toward glycolysis when stimulated with oxytocin. Additionally, myometrial cells rely on glucose utilization through glycolysis and glucose oxidation as their primary energy source and do not have the metabolic flexibility to utilize fatty acids when these pathways are inhibited.

## RESULTS

### Oxytocin exposure alters myometrial cell bioenergetic profiles

First, we aimed to determine the impact of short-term oxytocin exposure on mitochondrial function in the myometrial cell line hTERT-HM and in primary human myometrial smooth muscle cells (hMSMCs) obtained at planned (non-laboring) cesarean deliveries in women at term. We treated hTERT-HM cells and hMSMCs for one hour with oxytocin or vehicle and used the Seahorse XF Cell Mito Stress Test to measure oxygen consumption rate (OCR), extracellular acidification rate (ECAR), and associated bioenergetic parameters (Figure 1). Analysis of OCR (Figure 2A-B) showed that a 1-hour oxytocin treatment did not alter basal respiration rate, ATP linked respiration, or maximal respiration in either hTERT-HM cells (Figure 2C-D, G) or primary hMSMCs (Figure 2E-F, I). In contrast, spare capacity was significantly increased in hTERT-HM cells treated with 10^−8^ M and 10^−7^ M oxytocin (Figure 2H) and in primary hMSMCs treated with 10^−8^ M oxytocin (Figure 2J). A similar concentration-dependent increase in spare respiratory capacity was observed with acute oxytocin exposure in hTERT-HM cells at 10^−8^ - 10^−6^ M, but not in primary hMSMCs (Supplementary Figure S2).

**Figure 1:**
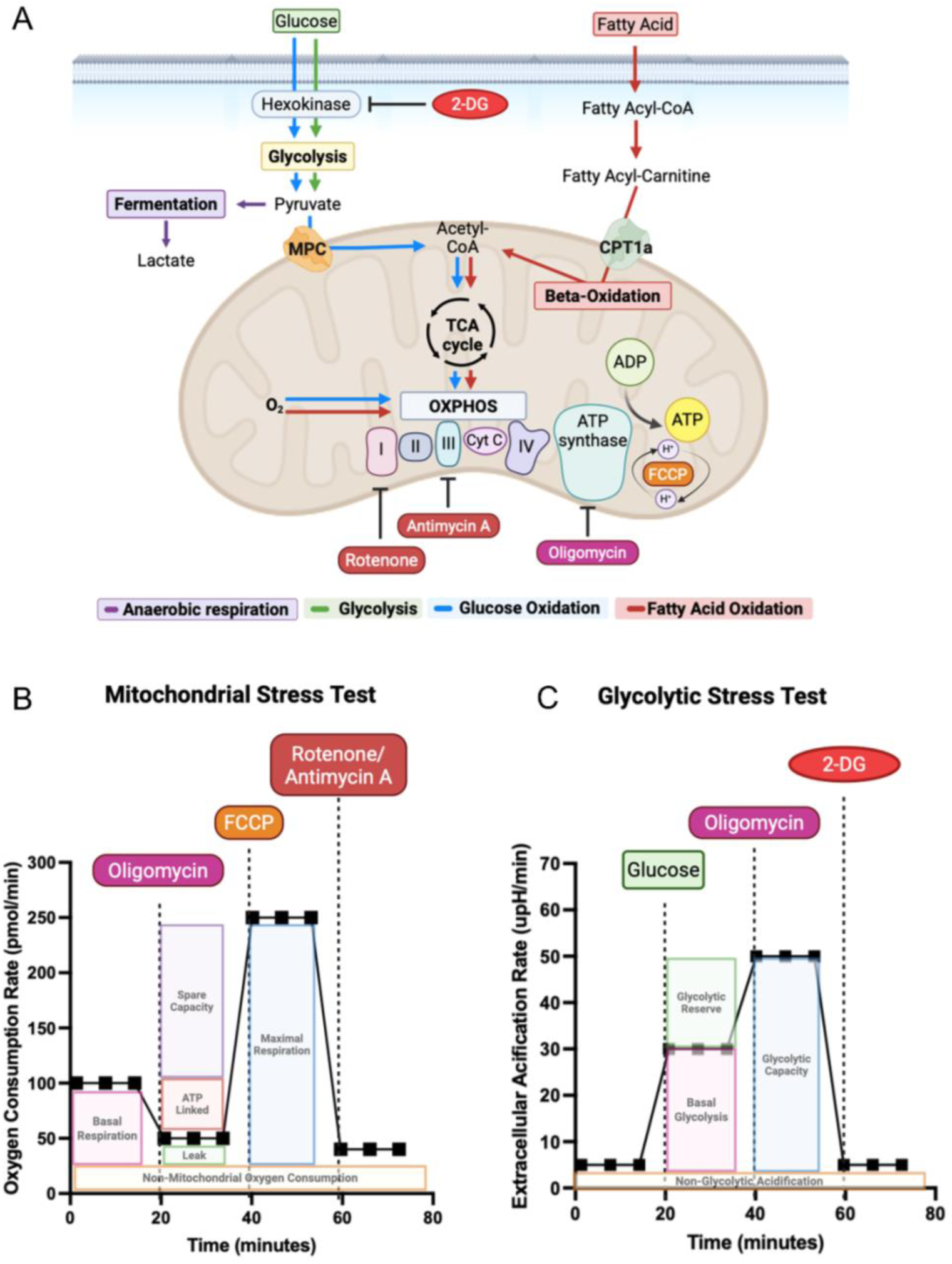
Experimental schematic for extracellular flux assay and the typical OCR and ECAR curves with Seahorse XF Mitochondrial Stress Test and Seahorse XF Glycolytic Stress Test assay parameters. (A) Metabolic pathways including glycolysis, electron transport chain, TCA cycle, OXPHOS. Anaerobic respiration is denoted in purple lines, glycolysis denoted in green lines, glucose oxidation denoted in blue lines, fatty acid beta oxidation denoted in red lines. (B) Seahorse XFe96 Analyzer records OCR values (y-axis) versus time (x-axis) before and after the injections of oligomycin, FCCP and rotenone/antimycin A. These OCR readings are used to calculate basal respiration, ATP linked respiration, maximal respiration, spare capacity, non-mitochondrial respiration, and proton leak. (C) Seahorse XFe96 Analyzer records ECAR values (y-axis) versus time (x-axis) before and after the injections of glucose, oligomycin, and 2-DG. These ECAR readings are used to calculate basal glycolysis, glycolytic reserve, glycolytic capacity and non-glycolytic acidification. OCR, oxygen consumption rate; TCA, tricarboxylic acid; ECAR, extracellular acidification rate; FCCP, trifluoromethoxy carbonyl cyanide phenylhydrazone; 2-DG, 2-deoxy-glucose; OXPHOS, oxidative phosphorylation.

**Figure 2.**
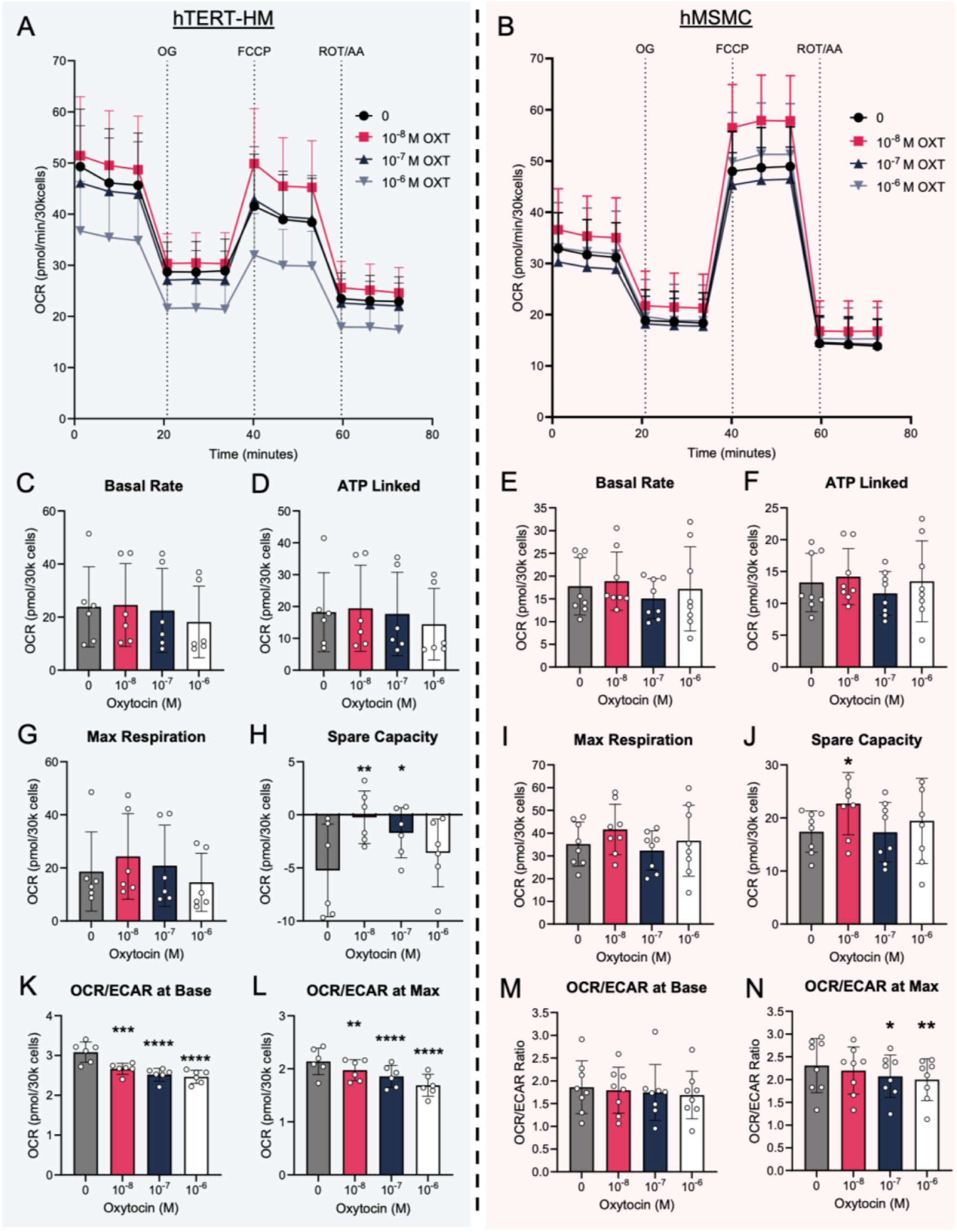
Short-term exposure to oxytocin increases mitochondrial spare capacity and decreases OCR:ECAR ratio. (A-B) Combined Mitochondrial Stress Test traces for hTERT-HM cells (n=6, blue) and hMSMCs (n=8, pink) treated for 1 hour with 10^−8^ M, 10^−7^ M, or 10^−6^ M oxytocin. Quantification of basal respiration (C, E), ATP linked respiration (D, F), maximal respiration (G, I), spare respiratory capacity (H, J), OCR:ECAR ratio at baseline (K, M) and OCR:ECAR ratio at maximal respiration (L, N). Results in A and B are presented as means +/− SEM. Results in C-N are presented as means +/− SD. \**P* < 0.05, \*\**P*< 0.01, \*\*\**P* < 0.001, \*\*\*\**P*< 0.0001 by repeated measures one-way ANOVA with Holm-Sidak multiple comparison test. OCR, oxygen consumption rate; ECAR, extracellular acidification rate; OXT, oxytocin; OG, oligomycin; FCCP, trifluoromethoxy carbonyl cyanide phenylhydrazone; ROT, rotenone; AA, antimycin A.

To further assess metabolic adaptation, we calculated the OCR/ECAR ratio as a measure of energy production via oxidative phosphorylation relative to glycolysis. At baseline, this ratio decreased in a concentration-dependent manner after 1-hour oxytocin treatment in both hTERT-HM and primary hMSMC cells (Figure 2K, M), suggesting a shift to glycolytic metabolism upon oxytocin treatment. This decrease persisted at maximal respiration in hTERT-HM cells when treated with 10^−8^ - 10^−6^ M oxytocin (Figure 2L) and in primary hMSMCs with 10^−7^ - 10^−6^ M oxytocin (Figure 2N). A similar concentration-dependent decrease in the OCR/ECAR ratio was observed with acute oxytocin exposure only in hMSMC cells at 10^−7^ - 10^−6^ M oxytocin (Supplementary Figure S2N).

Next, we examined whether these effects were sustained during prolonged oxytocin treatment. Because 10^−8^ M oxytocin had the greatest effect on mitochondrial bioenergetics in one-hour treatments, we used this dose for extended exposures. We treated hTERT-HM cells and primary hMSMCs with 10^−8^ M oxytocin and measured bioenergetic parameters at 2, 4, and 6 hours. Prolonged oxytocin exposure did not significantly alter basal respiration, ATP linked respiration, maximal respiration, or spare capacity in hTERT-HM cells or primary hMSMCs (Figure 3). However, prolonged oxytocin treatment consistently induced a decrease in the OCR/ECAR ratio in primary hMSMCs at baseline and maximal respiration (Figure 3M-N), indicating a metabolic shift toward glycolysis. These changes did not occur in hTERT-HM cells (Figure 3K-L). Thus, although oxytocin treatment only transiently increases spare capacity, the shift to glycolytic metabolism is persistent across all durations of oxytocin treatment in primary hMSMCs.

**Figure 3.**
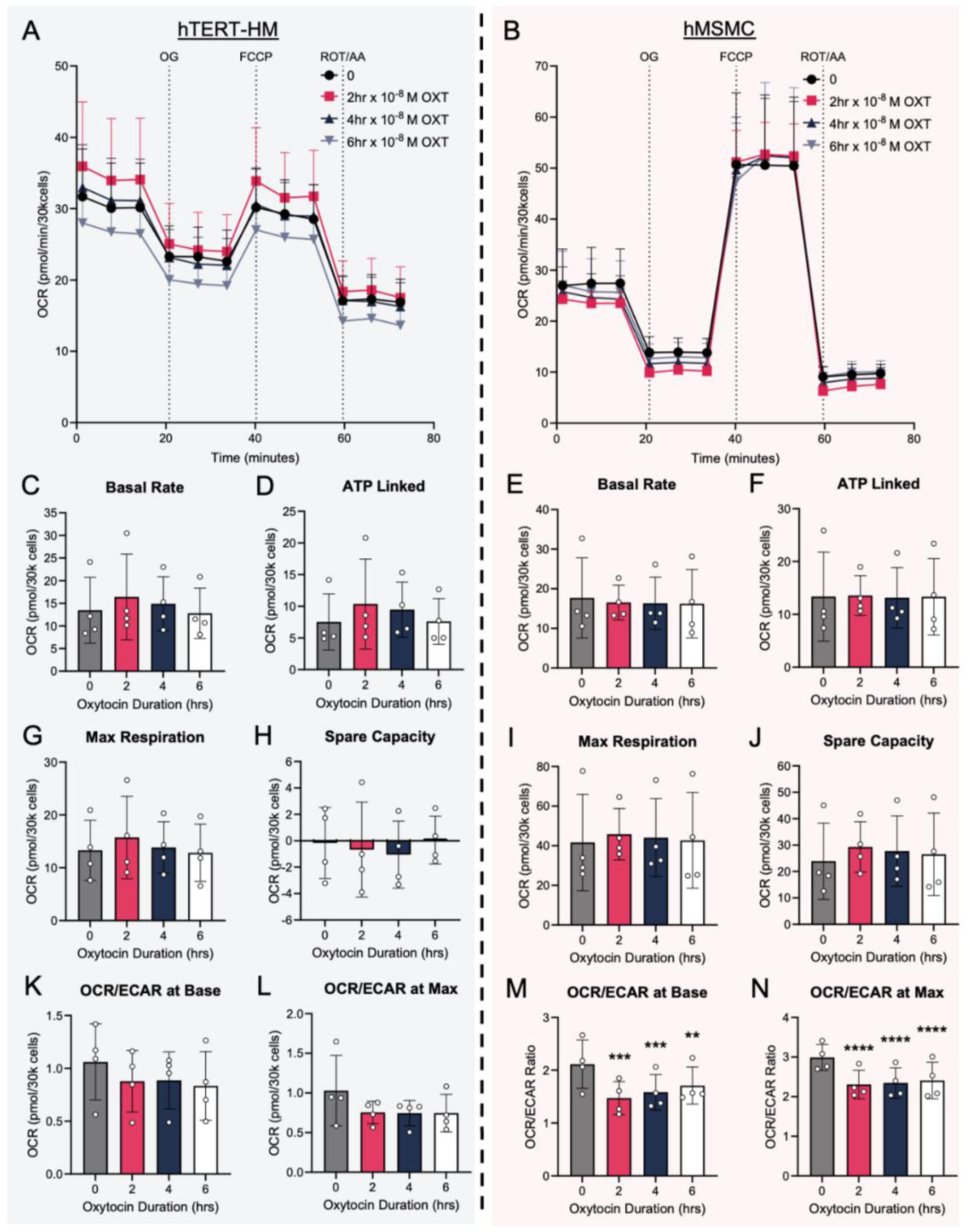
Prolonged oxytocin treatment does not affect mitochondrial spare capacity but decreases OCR:ECAR ratio. (A-B) Combined Mitochondrial Stress Test traces for hTERT-HM (n=4, blue) and hMSMC (n=4, pink) treated for 2, 4, 6 hours with 10^−8^ M oxytocin. Quantification of bioenergetic parameters, basal rate (C, E), ATP linked respiration (D, F), maximal respiration (G, I), spare respiratory capacity (H, J), OCR:ECAR ratio at baseline (K, M) and OCR:ECAR ratio at maximal respiration (L, N). Results in A and B are presented as means +/− SEM. Results in C-N are presented as means +/− SD. \**P* < 0.05, \*\**P*< 0.01, \*\*\**P* < 0.001, \*\*\*\**P*< 0.0001 by repeated measures one-way ANOVA with Holm-Sidak multiple comparison test. OCR, oxygen consumption rate; ECAR, extracellular acidification rate; OXT, oxytocin; OG, oligomycin; FCCP, trifluoromethoxy carbonyl cyanide phenylhydrazone; ROT, rotenone; AA, antimycin A.

### Oxytocin exposure alters myometrial cell glycolytic function

Given the alterations in spare respiratory capacity and OCR/ECAR ratios, we wondered whether oxytocin exposure affected energy production via the glycolytic pathway. To assess this, we used the Seahorse XF Glycolysis Stress Test to measure ECAR and glycolytic parameters. Acute treatment did not significantly increase the basal glycolytic rate or glycolytic reserve in either hTERT-HM cells or hMSMCs (Supplementary Figure S3). Glycolytic reserve was significantly decreased only with 10^−6^ M oxytocin in hTERT-HM cells (Supplementary Figure S3E). Similarly, there were no significant changes in glycolytic function following a one-hour oxytocin treatment of either cell line (Figure 4).

**Figure 4.**
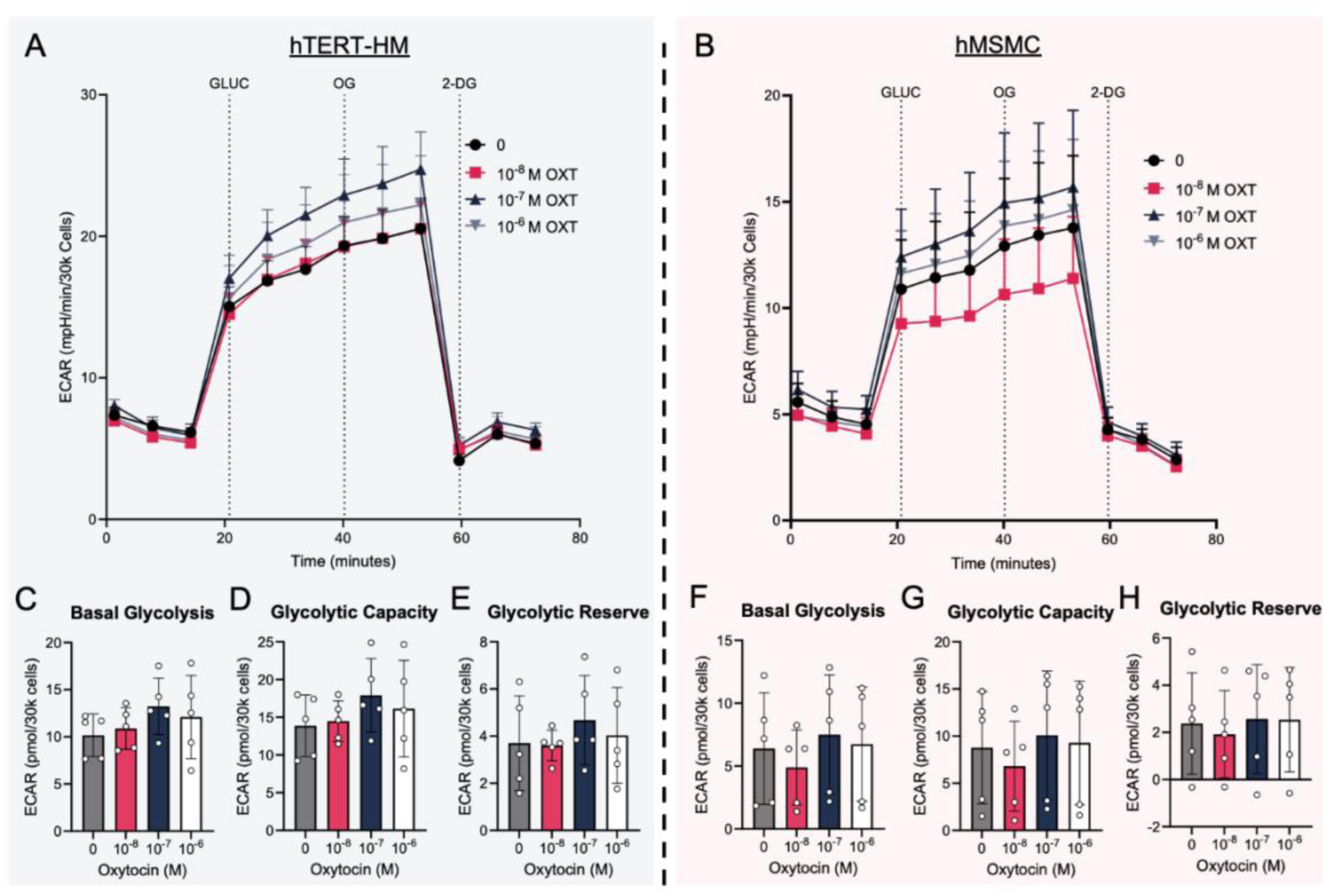
Short-term exposure to oxytocin does not significantly alter glycolytic function. (A-B) Combined Glycolytic Stress Test traces of hTERT-HM cells (n=5, blue) and hMSMCs (n=5, pink) treated for 1 hour with 10^−8^ M, 10^−7^ M, 10^−6^ M oxytocin. Quantification of bioenergetic parameters, basal glycolysis (C, F), glycolytic capacity (D, G), glycolytic reserve (E, H). Results in A and B are presented as means +/− SEM. Results in C-H are presented as means +/− SD. \**P* < 0.05, \*\**P*< 0.01, \*\*\**P* < 0.001, \*\*\*\**P*< 0.0001 by repeated measures one-way ANOVA with Holm-Sidak multiple comparison test. ECAR, extracellular acidification rate; OXT, oxytocin; GLUC, glucose; OG, oligomycin; 2-DG, 2-deoxy-glucose.

We then analyzed whether prolonged oxytocin treatment led to sustained changes in glycolytic function. hTERT-HM cells and hMSMCs were treated with 10^−8^ M oxytocin, and glycolytic parameters were analyzed after 2, 4, and 6 hours. Prolonged treatment with oxytocin significantly increased basal glycolysis and glycolytic capacity at 2 and 4 hours in hTERT-HM cells (Figure 5C-D). Similarly, in primary hMSMCs, basal glycolysis was significantly increased at 4 hours (Figure 5F), and glycolytic capacity was increased at 4 and 6 hours (Figure 5G). Glycolytic reserve was not altered by oxytocin in either hTERT-HM cells or hMSMCs (Figure 5E, H). Taken together, these results suggest that short-term oxytocin exposure has minimal effects on glycolytic function, whereas prolonged treatment enhances glycolytic activity by increasing basal glycolysis and glycolytic capacity in myometrial cells.

**Figure 5.**
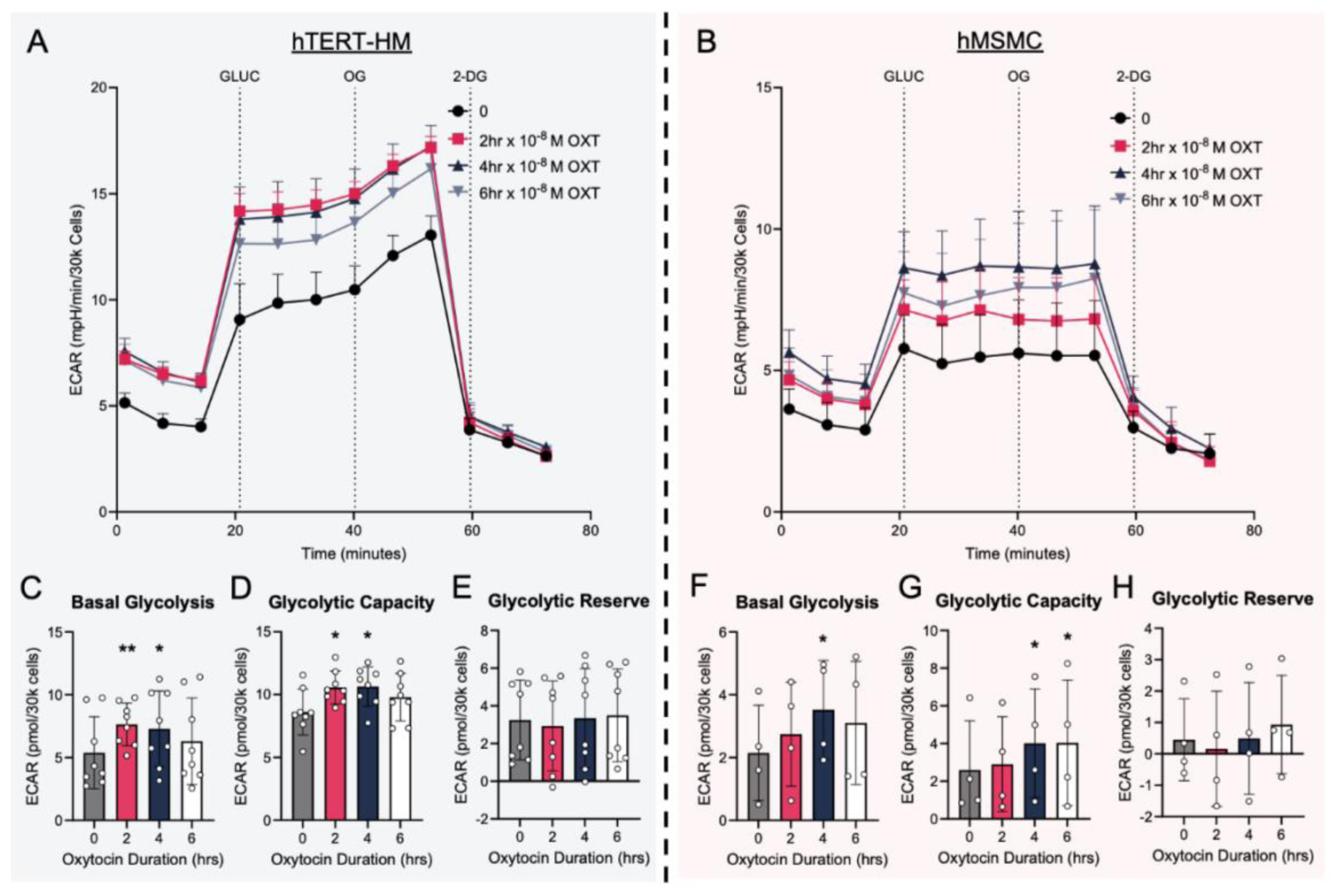
Prolonged oxytocin treatment increases basal glycolysis and glycolytic capacity. (A-B) Combined Glycolytic Stress Test traces for hTERT-HM cells (n=8, blue) and hMSMCs (n=4, pink) treated for 2, 4, 6 hours with 10^−8^ M oxytocin. Quantification of bioenergetic parameters, basal glycolysis (C, F), glycolytic capacity (D, G), glycolytic reserve (E, H). Results in A and B are presented as means +/− SEM. Results in C-H are presented as means +/− SD. \**P* < 0.05, \*\**P*< 0.01, \*\*\**P* < 0.001, \*\*\*\**P*< 0.0001 by repeated measures one-way ANOVA with Holm-Sidak multiple comparison test. ECAR, extracellular acidification rate; OXT, oxytocin; GLUC, glucose; OG, oligomycin; 2-DG, 2-deoxy-glucose.

### Myometrial cells preferentially use glucose as a main energy substrate

We next assessed whether myometrial cells preferentially use a specific substrate for energy production, whether they exhibit metabolic flexibility, and whether their substrate preferences are altered by exposure to oxytocin. Because hTERT-HM cells and hMSMCs exhibited similar responses in our earlier Mito Stress Test experiments, we only performed these experiments in primary hMSMCs. Bioenergetic parameters were measured in the Seahorse XF Substrate Oxidation Test, and specific inhibitors for each oxidation pathway were added individually or in combination (Figure 6 A-B). UK5099 was used to inhibit the mitochondrial pyruvate carrier and glucose oxidation, etomoxir was used to block carnitine palmitoyl transferase 1 and beta oxidation of fatty acids, and BPTES was used to block glutaminase 1 and glutamine oxidation.

**Figure 6.**
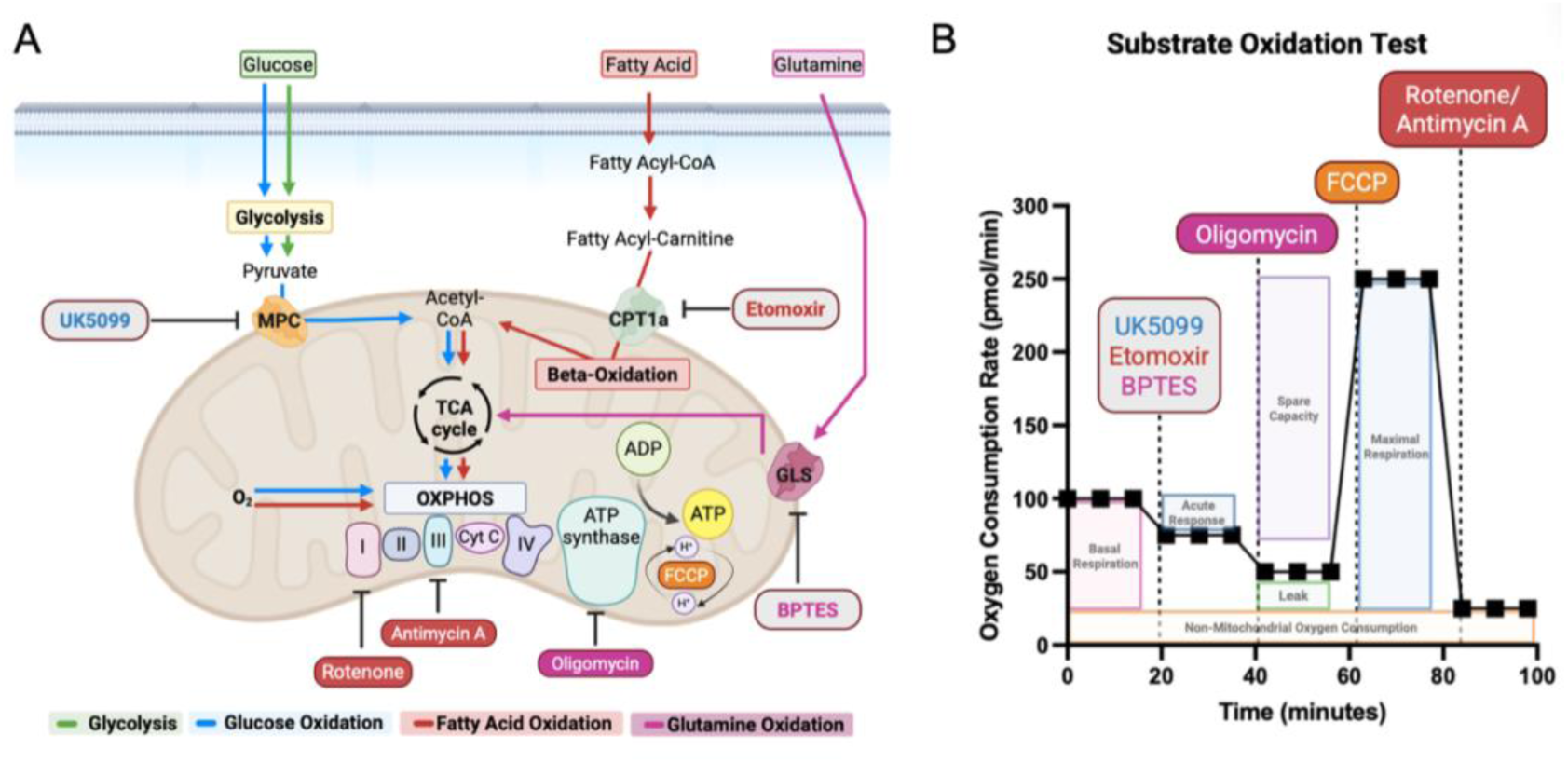
Experimental schematic for extracellular flux assay with the typical OCR curve of Substrate Oxidation Test assay parameters. (A) Metabolic pathways including glycolysis, electron transport chain, TCA cycle, OXPHOS. Glycolysis denoted in green lines, glucose oxidation denoted in blue lines, fatty acid beta oxidation denoted in red lines, glutamine oxidation denoted in magenta lines. (B) Seahorse XFe96 Analyzer records OCR values (y-axis) versus time (x-axis) before and after the injections of UK5099 and/or Etomoxir and/or BPTES, oligomycin, FCCP and rotenone/antimycin A. These OCR readings are used to calculate basal respiration, acute response to inhibitor(s), ATP linked respiration, maximal respiration, spare capacity, non-mitochondrial respiration, and proton leak. OCR, oxygen consumption rate; TCA, tricarboxylic acid; FCCP, trifluoromethoxy carbonyl cyanide phenylhydrazone; OXPHOS, oxidative phosphorylation; MPC, mitochondrial pyruvate carrier; CPT1a, carnitine palmitoyl transferase 1a; GLS, glutaminase.

Basal respiration significantly decreased only in the presence of UK5099, either alone or in combination, suggesting a strong reliance on glucose oxidation and limited metabolic flexibility (Figure 7A-D). Additionally, basal respiration in hMSMCs decreased in the presence of BPTES following oxytocin treatment, suggesting a contribution from glutamine oxidation under stimulated conditions (Figure 7D). In contrast, there was no effect on basal respiration following etomoxir treatment, indicating little reliance on fatty acid oxidation. In cells not exposed to oxytocin, only a combination of all three inhibitors significantly reduced maximal respiration (Figure 7E). However, following one-hour treatment with oxytocin, maximal respiration decreased in all treatment groups when glucose oxidation inhibitor UK5099 was present, whether alone or in combination (Figure 7F). Similarly, untreated hMSMCs had a significantly lower spare capacity in the presence of UK5099, but not etomoxir or BPTES alone (Figure 7G). This significant decrease persisted following the one-hour treatment with oxytocin (Figure 7H).

**Figure 7.**
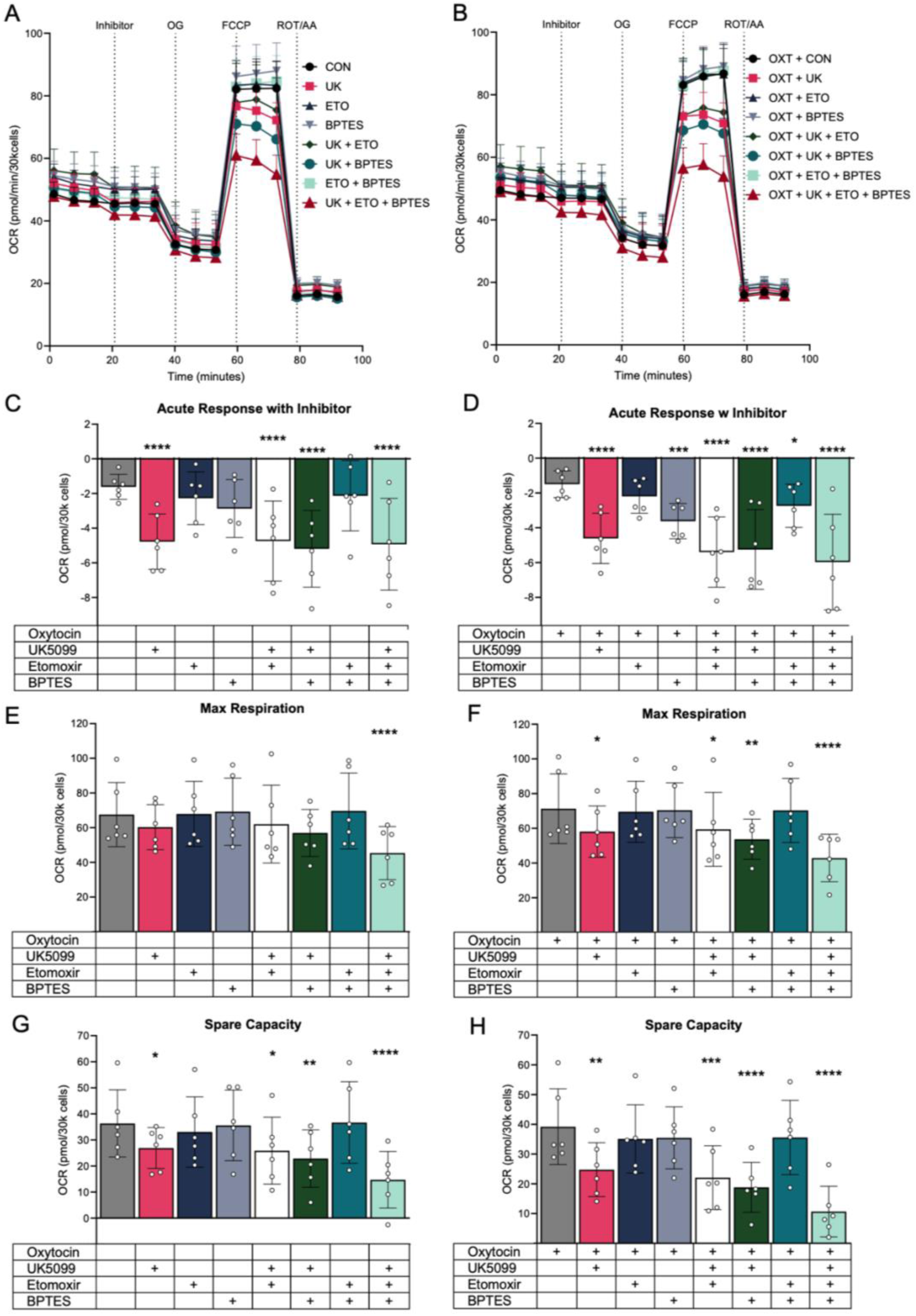
Myometrial cells preferentially use glucose as a main energy substrate. (A) Combined Substrate Oxidation Test traces of hMSMC untreated (n=6) or (B) treated for 1 hour with 10^−8^ M oxytocin (n=6). Substrate Oxidation Test was performed with inhibitors of oxidation pathways UK5099, etomoxir and BPTES individually or in combination to rigorously assess substrate preference. Quantification of bioenergetic parameters, acute response with inhibitor (C-D), maximal respiration (E-F) and spare respiratory capacity (G-H). Results in A and B are presented as means +/− SEM. Results in C-N are presented as means +/− SD. \**P* < 0.05, \*\**P*< 0.01, \*\*\**P* < 0.001, \*\*\*\**P*< 0.0001 by repeated measures one-way ANOVA with Holm-Sidak multiple comparison test. OCR, oxygen consumption rate; ECAR, extracellular acidification rate; OXT, oxytocin; UK, UK5099; ETO, etomoxir; OG, oligomycin; FCCP, trifluoromethoxy carbonyl cyanide phenylhydrazone; ROT, rotenone; AA, antimycin A.

Together, these results demonstrate that myometrial cells depend predominantly on glucose oxidation to support mitochondrial respiration and exhibit limited metabolic flexibility. Oxytocin treatment further heightens this glucose dependence under maximum respiration conditions.

### Oxytocin exposure modulates AMPK phosphorylation and LDH isoform expression

Given the observed shift towards glycolysis following oxytocin exposure, we next asked whether primary hMSMCs exhibited changes in phosphorylation of adenosine monophosphate-activated protein kinase (AMPK), a key regulator of cellular energy homeostasis. AMPK is activated via phosphorylation when the AMP/ATP ratio is increased, shifting intracellular metabolic pathways toward increased catabolism to restore energy balance. Conversely, decreased AMPK phosphorylation is associated with increased aerobic glycolysis. Western blot analysis showed that a one-hour treatment with 10^−7^ M oxytocin significantly decreased AMPK phosphorylation (Figure 8A-B). There were no differences in AMPK phosphorylation after longer treatments (Figure 8C-D).

**Figure 8.**
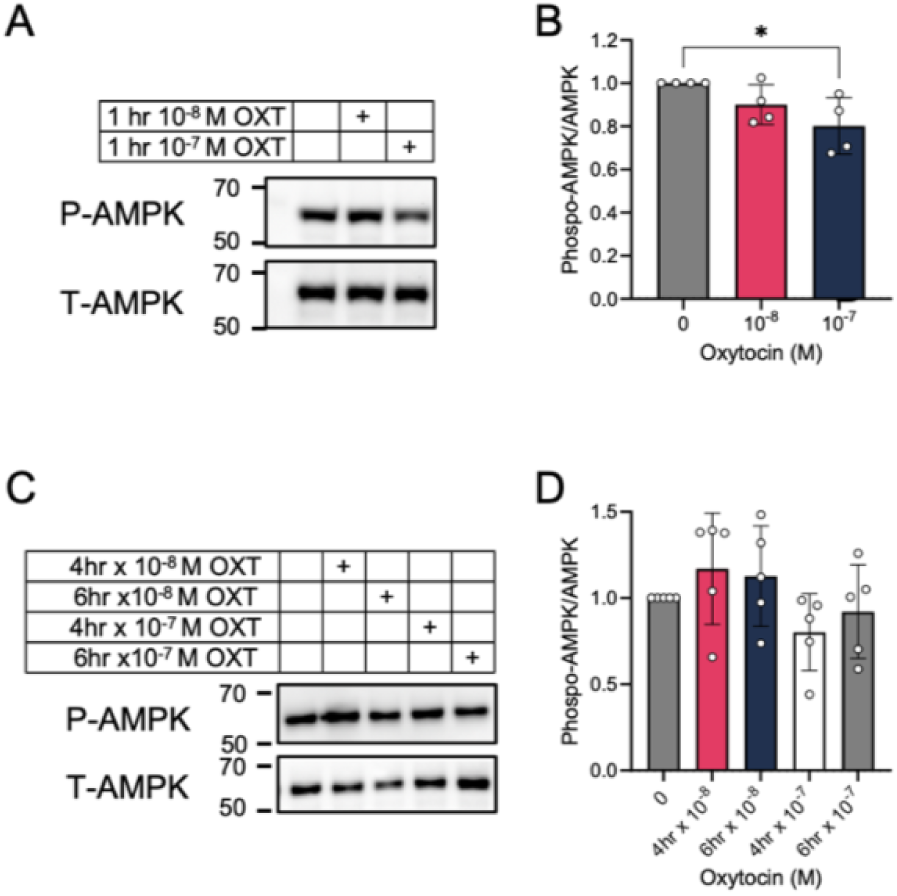
Short-term exposure to oxytocin decreases active AMPK. hMSMC cells were treated with (A) 10^−8^ M and 10^−7^ M oxytocin for 1 hour (n=4) or (C) 10^−8^ M and 10^−7^ M oxytocin for 4 and 6 hours (n=5). Western immunoblot with anti-P-AMPK antibody was used to quantitate protein and anti-T-AMPK was used as control. Representative immunoblots are shown. (B, D) Relative P-AMPK/T-AMPK, quantifications are shown. Results are presented as means +/− SD. \**P* < 0.05, \*\**P*< 0.01, \*\*\**P* < 0.001, \*\*\*\**P*< 0.0001 by repeated measures one-way ANOVA with Holm-Sidak multiple comparison test. P-AMPK, phosphorylated-AMPK; T-AMPK, total-AMPK.

To further assess glycolytic regulation, we quantitated lactate dehydrogenase subunits A (LDHA) and B (LDHB). LDHA converts pyruvate to lactate, whereas LDHB converts lactate to pyruvate. We observed no differences in LDHA or LDHB after one-hour treatment with either 10^−8^ M oxytocin or 10^−7^ M oxytocin (Figure 9A-E). However, after a four-hour treatment with 10^−7^ M oxytocin, LDHB abundance was significantly decreased (Figure 9G and I), but LDHA abundance was unchanged (Figure 9F and H). Furthermore, the LDHA/LDHB ratio (Figure 9J), a measure of lactate production versus oxidation, was significantly increased after a four-hour treatment with 10^−7^ M oxytocin, suggesting increased turnover of pyruvate to lactate via glycolysis. Together, these findings suggest that oxytocin transiently suppresses AMPK phosphorylation and shifts LDH isoform expression to favor lactate production, supporting the observed glycolytic reprogramming in myometrial cells.

**Figure 9.**
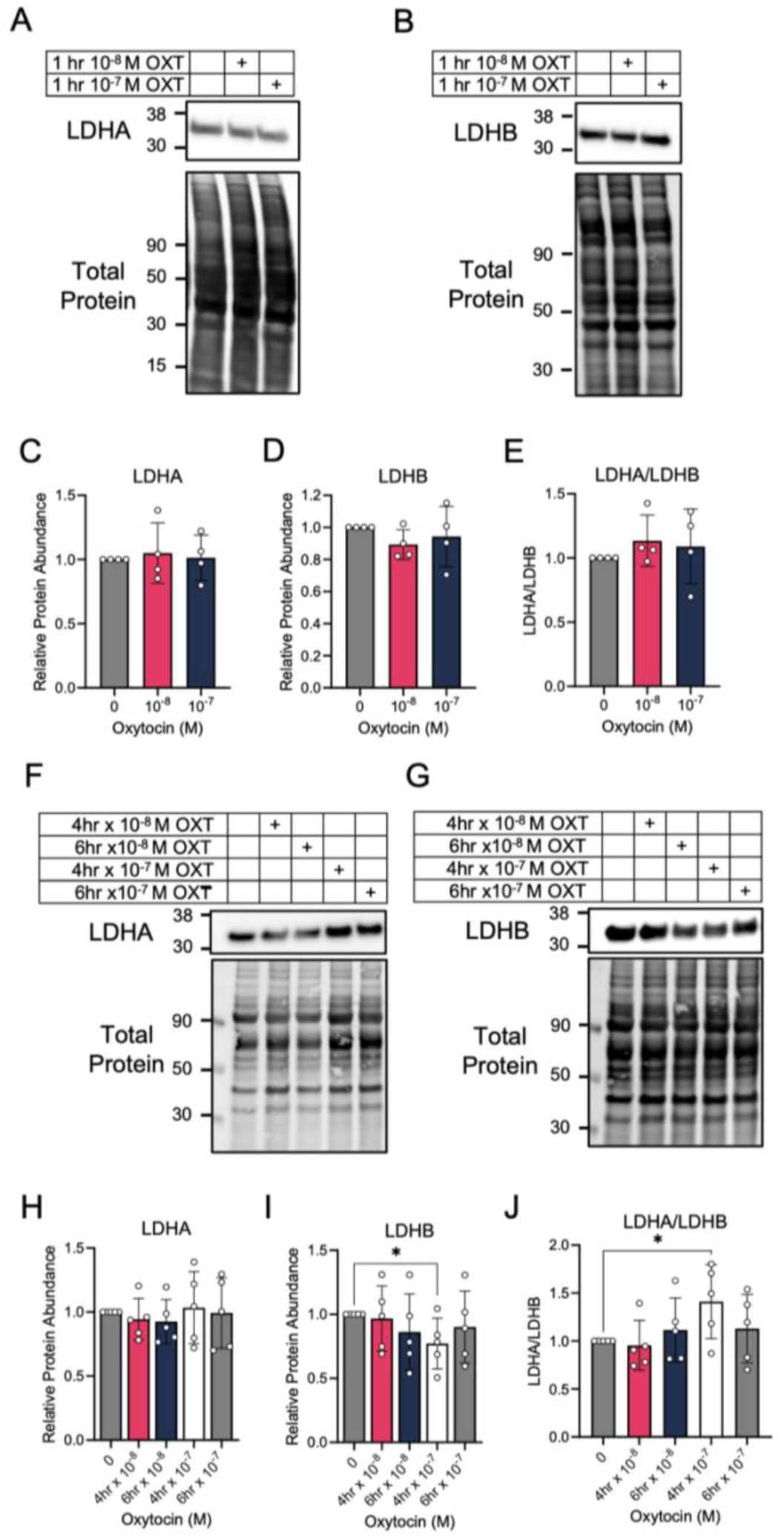
Prolonged oxytocin treatment decreases conversion of lactate to pyruvate and increases pyruvate to lactate turnover ratio. hMSMC cells were treated with (A-B) 10^−8^ M and 10^−7^ M oxytocin for 1 hour (n=4) or (F-G) 10^−8^ M and 10^−7^ M oxytocin for 4 and 6 hours (n=5). Western immunoblot with anti-LDHA (A, F) and anti-LDHB (B, G) antibodies were used to quantitate protein and total protein quantification was used as loading control. Representative immunoblots are shown. Relative LDHA (C, H), LDHB (D, I) and LDHA/LDHB (E, J) quantifications are shown. Results are presented as means +/− SD. \**P* < 0.05, \*\**P*< 0.01, \*\*\**P* < 0.001, \*\*\*\**P*< 0.0001 by repeated measures one-way ANOVA with Holm-Sidak multiple comparison test. LDHA, lactate dehydrogenase A; LDHB, lactate dehydrogenase B.

### Inhibition of glucose oxidation reduces myometrial contractility

Finally, we investigated whether modulating oxidation pathways *ex vivo* in myometrial strips would result in changes in contractility. We inhibited the two main energetic pathways, glucose and fatty acid oxidation, with sequential treatments of low and high doses of either UK5099 or etomoxir, in spontaneously contracting (Figure 10A-B) and oxytocin stimulated (Figure 10F-G) myometrial strips. Quantification of the area under the force-time curve (AUC), amplitude (AMP) and frequency (FREQ) showed that during spontaneous contractility myometrial strips had a significant decrease in fold change from baseline in AUC, AMP and FREQ (Figure 10C-E) when treated with 9 µM UK5099, but not etomoxir. Additionally, there was a significantly lower AUC (Figure 10C) in myometrial strips treated with 6 µM UK5099 compared to etomoxir, and AMP (Figure 10D) when treated with 9 µM UK5099 compared to etomoxir. In the setting of oxytocin stimulation, myometrial strips had a significant decrease in fold change from baseline in AUC (Figure 10H) when treated with 6 and 9 µM UK5099 and 9 µM etomoxir. FREQ (Figure 10J) was also significantly lower in the presence of 9 µM UK5099 but not etomoxir. Furthermore, there was a significantly lower AUC (Figure 10H) in myometrial strips treated with UK5099 compared to etomoxir at both doses, and significantly lower FREQ (Figure 10J) in strips treated with 9 µM UK5099 compared to etomoxir. Taken together, these findings suggests a reduction of myometrial contractility when glucose oxidation is inhibited supporting the observed reliance on glucose in myometrial cells *in vitro*.

**Figure 10.**
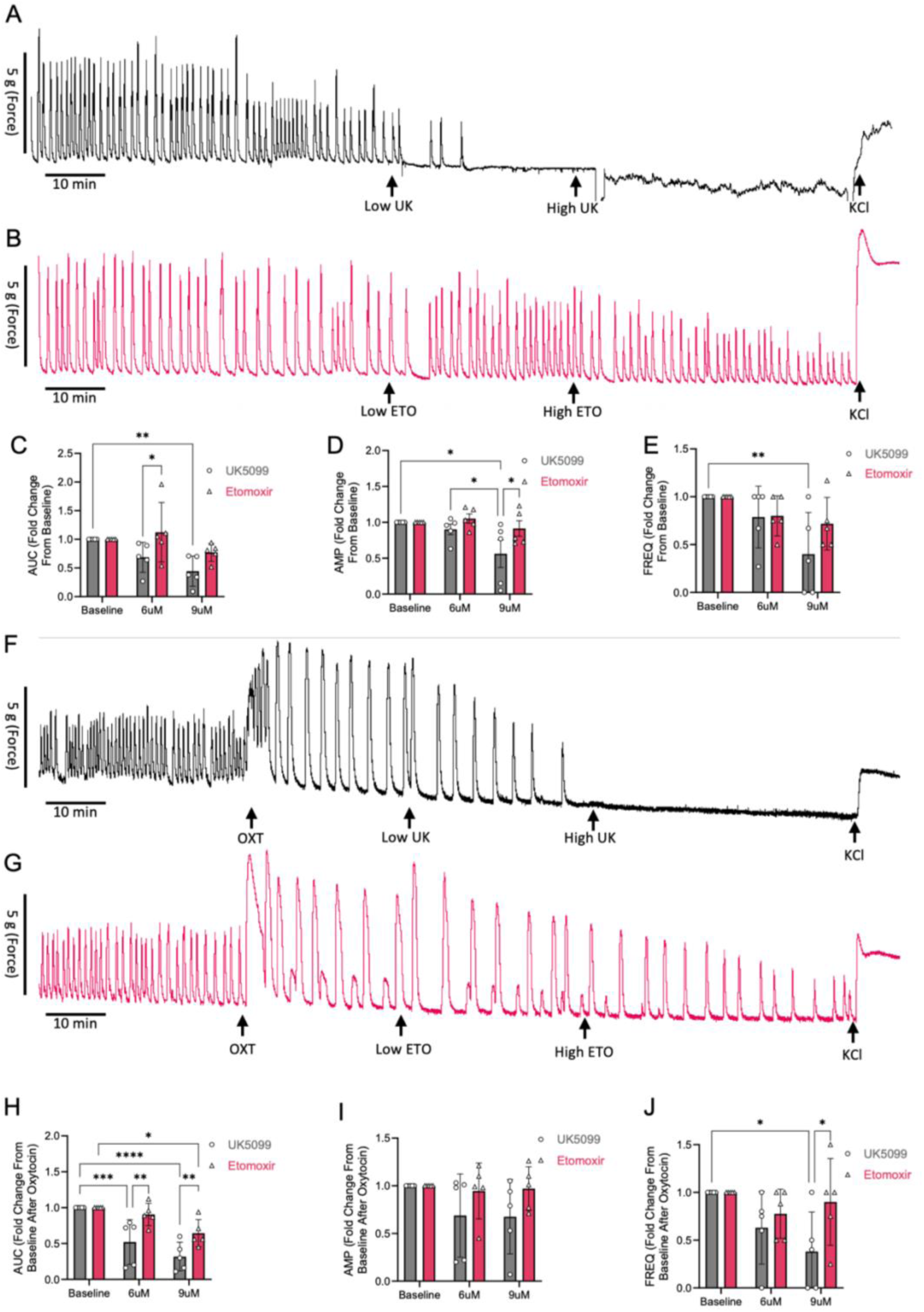
Inhibition of glucose oxidation decreases myometrial contractility. Representative *ex vivo* myometrial contractility traces from (A) spontaneous contractility with UK5099 (n=5, black), (B) spontaneous contractility with etomoxir (n=5, pink), (F) oxytocin stimulated contractility with UK5099 (n=5, black), (G) oxytocin stimulated contractility with etomoxir (n=5, pink). Area under the curve (C, H), amplitude (D, I) and frequency (E, J) was quantified as fold change from individual channel baseline. Results are presented as means +/− SD. \**P* < 0.05, \*\**P*< 0.01, \*\*\**P* < 0.001, \*\*\*\**P*< 0.0001 by two-way ANOVA with Holm-Sidak multiple comparison test. OXT, oxytocin; UK, UK5099; ETO, etomoxir; AUC, area under the curve; AMP, amplitude; FREQ, frequency.

## DISCUSSION

Here, we have shown that myometrial cells primarily rely on oxidative phosphorylation during quiescence and shift their energy production toward glycolysis when stimulated with oxytocin. Additionally, spare respiratory capacity in the mitochondria and cellular glycolytic capacity increase after treatment with oxytocin, suggesting that myometrial cells can adapt their energy use in states of higher energy demands. The decreased basal OCR upon treatment with UK5099, but not etomoxir or BPTES, suggests a preference for glucose oxidation under basal conditions in quiescent myometrial cells. This is also supported by the reduction in myometrial contractility upon treatment with UK5099, but not etomoxir. Further, the additional decrease in basal OCR upon BPTES treatment, but not etomoxir, following one-hour treatment with oxytocin indicates that myometrial cells can shift from glucose to glutamine but not to fatty acid oxidation under conditions of higher metabolic demand. Finally, the reduced mitochondrial spare respiratory capacity after treatment with UK5099 in the non-exposed and oxytocin-exposed cells further demonstrates reliance on glucose oxidation in quiescent and contractile myometrial cells.

Our data are consistent with previous research suggesting that, under normal metabolic conditions, the myometrium at term preferentially uses glucose as its energy substrate^2–5^. Additionally, the myometrium appears to use most of this glucose via anaerobic respiration even in the presence of normal oxygen concentrations, generating lactate in the process^3,15^. Furthermore, a previous study of bovine myometrium sugested that escalating concentrations of fatty axid oxidation inhibitor etomoxir increased myometrial contractility likely due to an increase in glucose utilization^16^. These observations are consistent with our findings that inhibition of glucose oxidation via addition of UK5099 decreased the OCR and myometrial contractility and that oxytocin-stimulated myometrial cells have an increased glycolytic capacity as measured by the ECAR. Furthermore, the significant decrease in the OCR/ECAR ratio at maximal respiration after oxytocin treatment suggests that myometrial cells readily switch to utilization of glucose via glycolysis after ATP synthase is inhibited. Additionally, this is also consistent with the reduction of spontaneous and oxytocin-induced myometrial contractility with inhibition of glucose oxidation by UK5099.

In contrast, Qian et al. showed that laboring human myometrium contained more metabolites associated with fatty acid oxidation and lipolysis than did nonlaboring myometrium, suggesting that utilization of fatty acids increases during labor^17^. Myometrial cells in our study did not readily switch to fatty acids, as evidenced by no additional change in OCR when etomoxir was added to UK5099 treatment. Additionally, we saw a significant decrease in all contractility parameters with both low and high doses of UK5099 treatment in both spontaneous and oxytocin-induced contractility, while the only change with etomoxir was a decrease in contraction AUC in oxytocin-stimulated myometrial strips following high dose etomoxir exposure. This may reflect a small reliance on fatty acid utilization under high energy demand conditions, but could also be due to etomoxir having off target and non-specific effects in other cell types in the uterine strips^18,19^. Either way, the etomoxir effect was minimal compared to inhibition of glucose oxidation by with UK5099. It is important to note that *in vitro* we used primary myometrial cells from women who were at term gestation but were not laboring and that we modelled contractility by treating the cells with oxytocin. This strategy likely did not fully recapitulate the myometrial laboring contractile phenotype. Further work is needed to determine the energetic profiles and metabolic flexibility of myometrial cells collected during active labor. However, this difference may partially explain the higher cesarean rates in cases of labor induction than in spontaneous labor. Furthermore, myometrial energy utilization may be a viable target to improve labor outcomes.

In states of altered substrate availability, such as obesity, excess aggregation of fatty acids in skeletal and cardiac muscle results in increased fatty acid β-oxidation and inhibition of glucose oxidation. The resulting failure to metabolically switch substrates causes contractility dysregulation^6,20,21^. We previously showed that maternal diet-induced obesity in mice is associated with upregulated long-chain fatty acid catabolism and accumulation of medium-chain fatty acids in the uterus^22^. In the future, mitochondrial bioenergetics experiments could be performed to determine substrate preferences and potential for metabolic flexibility in myometrial cells from women with obesity.

A surprising finding was the decrease in basal respiration in oxytocin-treated cells when BPTES was present, suggesting that glutamine oxidation supplements glucose oxidation when metabolic demand increases. However, BPTES did not affect maximal respiration or spare capacity, implying that, at peak energy production, glucose remains the dominant substrate. Glutamine is used as an energy source in many cell types including myocytes^23–25^. When the energy demand in skeletal muscle increases, such as during prolonged exercise, glutamine becomes essential for increasing the tissue strength and performance^26^. Whereas muscle cells rely heavily on oxidative phosphorylation for ATP generation^27–30^, other cell types, such as cancer cells, utilize anaerobic respiration even in the presence of oxygen. This phenomenon, known as the Warburg effect^31^, occurs when cells depend on glutamine metabolism for the high energy demands of fast proliferation^32^. Glutamine also plays a role in cell proliferation in vascular smooth muscle cells, inducing a switch towards a more synthetic and less contractile phenotype *in vitro*^33^. Although further studies are needed to properly characterize the complex interactions between glucose and glutamine metabolism, which vary between cell types, these substrates exhibit compensatory effects to maintain homeostasis within the TCA cycle^25,34^. It is possible that, in myometrial cells stimulated with oxytocin, glutamine oxidation is coupled to glucose oxidation to adapt to higher energy demands at baseline, as shown by the sensitivity to both UK5099 and BPTES. However, the unchanged maximal respiration and spare capacity with BPTES treatment suggests that, once maximal respiration is engaged, contribution to ATP production from glutamine oxidation becomes negligible and glucose remains the primary energy source.

Our metabolic data indicate that a short exposure to oxytocin (≤1 hour) significantly increases spare respiratory capacity, which returns to baseline with prolonged exposure to oxytocin (≥2 hours). This suggests that myometrial cells can increase energy production to adapt to higher energy demands but cannot sustain this level of oxidative metabolism for a prolonged period. The decrease in OCR/ECAR ratio we observed starting at one hour persisted with oxytocin treatment at two, four, and six hours and is indicative of decreased oxidative phosphorylation and increased glycolysis. Additionally, basal glycolysis and glycolytic capacity were significantly increased after two hours of oxytocin stimulation, suggesting that myometrial cells switch to glycolysis to maintain ATP production for contractility even when oxygen levels are not restricted. It is possible that spare capacity returned to baseline after prolonged exposure to oxytocin because of oxytocin receptor desensitization and internalization^35,36^. However, this is unlikely given the concurrent increase in glycolysis at these timepoints.

The functional significance of aerobic glycolysis in smooth muscle is unclear, but research suggests it is coupled to ion channel activity in the myometrium^37^. Decreased oxygenation lowers Ca^2+^ and excitability, increasing K^+^ channel activity, hyperpolarization, and inhibition of depolarizing ion channels^38,39^. ATP-gated potassium channels (K_ATP_), which are abundant in the myometrium, link metabolic changes to membrane excitability^40–42^. When ATP concentrations are low, K_ATP_ channels open, resulting in an efflux of K^+^ and hyperpolarization. When ATP concentrations are high, K_ATP_ channels close, depolarizing the membrane and allowing an influx of Ca^2+^ via voltage-gated Ca^2+^ channels^41^. K_ATP_ channels in vascular smooth muscle also respond to low pH, opening to promote vasodilation and requiring higher ATP concentrations to close^43,44^. Here, we show an increase in extracellular acidification with oxytocin stimulation, suggesting that glycolysis-driven pH change could keep K_ATP_ channels open to maintain ATP during high energy demand. Previously, we found that maternal diet-induced obesity reduces K_ATP_ activity and associated contractile frequency in term non-laboring mouse myometrium^45^. Future studies are needed to assess K_ATP_ function during normal labor in response to agonist stimulation or in low pH conditions. Such work could reveal strategies to enhance myometrial contractility.

One limitation of this study is that isolated myometrial cells may behave differently than uterine tissue, which also contains stromal and innate immune cells^46^. Additionally, the isolated myometrial cells were not under tension. Although other smooth muscles spontaneously contract in response to increased tension, uterine contractility is suppressed during fetal growth^47^. However, mechanical stretch can induce changes in the myometrium^48,49^. For example, myometrial stretch can increase the expression of gap junction and agonist receptor genes that are required for labor initiation^48^. Despite the strong forces coming from the growing fetus and placenta, these genes are not activated until term^49^. During pregnancy, many mechanosensitive channels in the myometrium regulate membrane polarization, such as the Ca^2+^ channel TRPV4^50^ and PIEZO1^51,52^. These stretch-activated channels play an important role in mediating quiescence and contractility. Whether stretch directly affects myometrial metabolism remains to be explored.

Finally, although we used oxytocin to mimic myometrial contractility, it did not model hypoxia^38^. In the myometrium during labor, the force generated by contractions compresses the spiral arteries and causes a transient ischemic and hypoxic event, resulting in a metabolic switch from aerobic to anaerobic respiration^53^. This was shown in an *in vivo* rat model, in which decreased blood flow and hypoxia led to decreased pH and increased lactate production resulting from anaerobic respiration^53^. Prolonged hypoxia decreases contractility in both murine and human myometrium^15^. This persists even in the presence of oxytocin^54,55^, suggesting that treatment with an agonist can dampen the decrease in contractility but not fully prevent it. In these studies, hypoxia or chemical metabolic inhibition occurred as a single event lasting for the entirety of the experimental procedure. In labor, hypoxia occurs at a higher frequency for a shorter duration^38^. Alotaibi et al. found that, when rat myometrium was exposed to short-term hypoxia, contractility increased. They termed this phenomenon hypoxia-induced force increase^56^. In the presence of oxytocin, hypoxia-induced force increase further augmented myometrial contractility^56^. How intermittent hypoxia alters the myometrial metabolic demand and preference is an important question to answer, particularly given this tissue’s unique preference for anaerobic respiration.

In conclusion, our data show that quiescent myometrial cells generate energy primarily via oxidative phosphorylation, whereas oxytocin-stimulated myometrial cells preferentially utilize glycolysis for energy production. Following oxytocin treatment, metabolic demand increases, and myometrial cells adapt their energy utilization by increasing their respiratory spare capacity and glycolytic capacity. Additionally, myometrial cells have a strong preference for glucose as a main source of energy and myometrial contractility is decreased when glucose oxidation is inhibited.

## MATERIALS AND METHODS

### Cell culture

Immortalized human myometrial cells (hTERT-HM)^57^ were maintained in Dulbecco’s modified Eagle’s/Ham’s F12 medium (DMEM; Gibco) without phenol red and supplemented with 10% fetal bovine serum (FBS; Gibco) and 25 μg/mL gentamicin (Gibco). Cells were kept in a humidified cell culture incubator at 37°C with 5% CO_2_. Oxytocin (Tocris) was utilized to stimulate myometrial contractility at different concentrations (10^−8^ M, 10^−7^ M, 10^−6^ M) and time points 1, 2, 4, and 6-hours.

### Isolation of primary human myometrial smooth muscle cells

Primary human myometrial smooth muscle cells (hMSMCs) were isolated from biopsies collected from the upper aspect of the hysterotomy performed at the time of term scheduled (unlabored) cesarean delivery. Deidentified human myometrial tissue was procured through the Washington University Reproductive Specimen Processing and Banking core. Within two hours of collection, the myometrial tissue was washed in ice-cold sterile Hanks’ Balanced Salt Solution (Gibco) supplemented with 5 μg/mL amphotericin B (Corning) and 50 μg/mL gentamicin. Tissue was cut into 1-2 mm pieces and incubated under constant rotation at 37 °C for 60 min in digestion solution: 1 mg/mL collagenase 1A and XI (Sigma) with gentamicin diluted in 4 mL DMEM with 1% charcoal stripped FBS (Gibco). The solution was then filtered through a 100 μM sterile cell strainer (CellTreat Scientific Products). Collagenase activity was stopped by adding 20 mL of warm 10% FBS + DMEM through the cell strainer. The cell suspension was centrifuged for 5 minutes and the supernatant was discarded. The cell pellet was resuspended in 3 ml of ACK Lysis Buffer (Gibco). After 5 minutes, 3 mL of 10% FCS + DMEM was added, the cell suspension was centrifuged again, and the supernatant discarded. Cells were plated and cultured in DMEM supplemented with smooth muscle cell growth media 2 mix (Promo Cell) and 50 μg/mL gentamicin. All cells were frozen at P1 and thawed to use as needed. Once thawed, 30,000 hMSMCs in passage 2-4 were plated on Seahorse 96-well culture plates (Agilent).

### Preparation of cells for bioenergetic analysis

To prepare Mitochondrial Stress Test and Substrate Oxidation Test medium, Seahorse XF DMEM medium, pH 7.4 (Agilent), was supplemented with 1 mM sodium pyruvate (Corning), 2 mM glutamine (Corning), and 10 mM glucose (Sigma). To prepare Glycolysis Stress Test medium, Seahorse XF DMEM medium, pH 7.4 (Agilent), was supplemented with 2 mM glutamine (Corning). Cell culture growth medium was removed from cells plated in Seahorse 96-well culture plates, and cells were then washed twice with pre-warmed assay medium to remove bicarbonate. After washing, cells were placed in 180 μL of assay medium, and the cell culture plate was placed in a 37 °C non-CO_2_ incubator for 45–60 min until loading it into the Seahorse XFe96 Analyzer (Agilent).

### Extracellular flux analysis

One day before the experiment, a Seahorse XFe96 sensor cartridge (Agilent) was hydrated by filling each well with 200 uL of calibration solution (Agilent), sensors were submerged into it, and it was placed in a non-CO_2_ incubator at 37 °C overnight. On the day of the experiment, the sensor cartridge was taken out of the non-CO_2_ incubator, and the injection ports were filled with appropriate compounds for each assay. For the Mitochondrial Stress Test, the following compounds were used: oligomycin (OG, ATP synthase complex V inhibitor), trifluoromethoxy carbonyl cyanide phenylhydrazone (FCCP, uncoupling agent that abolishes the proton gradient), and a combination of rotenone (ROT, complex I inhibitor) with antimycin A (AA, complex III inhibitor) to interrupt mitochondrial respiration. For the Substrate Oxidation Test, the following compounds were used: UK5099 (glucose and pyruvate oxidation inhibitor), etomoxir (ETO, long chain fatty acid oxidation inhibitor), BPTES (glutamine oxidation inhibitor) followed by OG, FCCP and ROT/AA. For the Glycolysis Stress Test, the following compounds were used: glucose (saturating dose fueling glycolysis), oligomycin (mitochondrial ATP synthase complex V inhibitor, which increases reliance on glycolysis), 2-deoxy-glucose (2-DG, competitive inhibitor of glucose, which inhibits glycolysis). Compounds were reconstituted, aliquoted, and store at −80 °C. On the day of the assay, an aliquot was thawed and used to prepare the stock solutions in assay medium and then further diluted to 10X concentrations before being loaded into the ports. The following final concentrations were used: 1.5 μM OG (Cayman Chemical), 0.5 μM ROT (Sigma), 0.5 μM AA (Cayman Chemical), 2.0 μM UK5099 (Selleckchem), 4.0 μM Etomoxir (Tocris), 3.0 μM BPTES (Cayman Chemical), 10 mM glucose (Sigma), and 50 mM 2-DG (Sigma).

Because the optimal FCCP concentration varies according to the metabolic demands of cells and can be inhibitory at high concentrations^58^, initial FCCP titration (0.25, 0.5, 1.0, 2.0, 4.0 μM) experiments were performed to identify the optimal concentration yielding the maximal oxidative capacity for hTERT-HM cells and primary hMSMCs. These concentrations were 0.5 μM and 4.0 μM, respectively (Supplementary Figure S1). Additionally, the oligomycin concentration (1.5, 2.5 μM) was titrated according to the manufacturer’s instructions to identify the optimal concentration for ATP synthase complex V inhibition for hTERT-HM cells and primary hMSMCs. The optimal concentration was 1.5 μM for both (Supplementary Figure S1). The loading volumes of each 10X compounds into the injection ports were 20 μL for Port A, 22 μL for Port B, 25 μL for Port C, and 27 μL for Port D. When the calibration process was complete (after about 20 min), the utility plate was replaced with the culture plate. After a short mixing period, the first set of measurements were recorded under basal conditions. Three measurements 8 min apart of oxygen consumption rate (OCR) and extracellular acidification rate (ECAR) were recorded between each injection.

### Calculation of metabolic parameters for mitochondrial function

Mitochondrial stress test (Figure 1B) and substrate oxidation test (Figure 6B) analysis provided direct measurements of basal respiration, ATP linked respiration, maximal respiration, spare capacity, non-mitochondrial respiration, and proton leak. Figure 1B shows a typical OCR curve and the mitochondrial stress test assay parameters analyzed. These metabolic parameters were calculated for each well by using the OCR plot as outlined in Figure 1B. Averages of individual well results (n = 3-4) for each sample were then calculated. Because cells were counted before the assay and equal numbers of cells were plated per well for each sample, Seahorse data were internally normalized to cell count. ECAR is driven by lactate accumulation in the extracellular medium. For the mitochondrial stress test assay, a proportional correlation between ECAR and lactate production was assumed, and changes in ECAR were used to indirectly reflect glycolytic capacity. Estimations of energy generation via oxidative phosphorylation vs. glycolysis were calculated at the basal rate of acidification (OCR/ECAR at base), after adding oligomycin to inhibt ATP-linked respiration (OCR/ECAR after OG), and after adding FCCP to induce maximal ATP requirement (OCR/ECAR at max).

### Calculation of metabolic parameters for glycolytic function

The glycolysis stress test (Figure 1C) provided direct measurements of basal glycolysis, glycolytic reserve, glycolytic capacity, and non-glycolytic acidification. Figure 1C shows a typical ECAR curve and the glycolysis stress test assay parameters analyzed. These metabolic parameters were calculated for each well by using the ECAR plot as explained in Figure 1C. Averages of individual well results (n = 3-4) for each sample were then calculated. Because cells were counted before the assay and equal numbers of cells were plated per well for each sample, Seahorse data were internally normalized to cell count.

### Western blot analyses

Cells were resuspended in lysis buffer (50 mM Tris HCl [pH 8.0], 150 mM NaCl, 1 mM EDTA, 0.5% Nonidet P-40, 2% Glycerol, complete Protease Inhibitor Cocktail [Roche Applied Science], and Phosphatase Inhibitor Mini Tablets [ThermoFisher]). Lysates were centrifuged at 14,000 g for 10 min at 4 °C. Protein concentration was quantified with the BCA Protein Assay Kit (Pierce). Proteins were then separated by SDS-PAGE in 4-12% Bolt Bis-Tris gels (ThermoFisher) and transferred to Immobilon-FL PVDF Membrane (Millipore Sigma). Membranes were stained for total protein with REVERT™ Kit (LI-COR) according to the manufacturer’s instructions, and total protein was used as the loading control. Membranes were then blocked in 1x Blocker FL Fluorescent Blocking Buffer (ThermoFisher) for 1 hour. Primary and secondary antibodies were diluted in 1x Blocker FL Fluorescent Blocking Buffer (ThermoFisher). Membranes were incubated with primary antibodies against the following proteins: LDHA (1:1000, Proteintech Cat# 19987-1-AP, RRID: AB_10646429), LDHB (1:1000, Proteintech Cat# 14824-1-AP, RRID: AB_2134953), phosphorylated AMPKα Thr172 (1:1000, Cell Signaling Technology Cat# 2535, RRID: AB_331250), or total AMPKα (1:1000, Cell Signaling Technology Cat# 2532, RRID: AB_330331). They were then incubated with anti-rabbit IgG (H+L) (1:10,000 Jackson ImmunoResearch Labs Cat# 111-035-003, RRID: AB_2313567) HRP-conjugated secondary antibody. The signal was detected with Super Signal West Femto Maximum Sensitivity Substrate (ThermoFisher). Membranes were imaged on a ChemiDoc MP (Bio-Rad Laboratories) and quantitated in Image Lab (Bio-Rad Laboratories). Values for LDHA and LDHB proteins were normalized to total protein. Phosphorylated AMPKα was normalized to total AMPKα.

### Ex vivo contractility

Myometrium was collected from patients at time of scheduled c-section and dissected into 8 x 2mm strips. The strips were placed in an organ bath warmed to 37°C and gassed with 95% O_2_ and 5% CO_2_ and immersed in physiologic Krebs buffer (133 mM NaCl, 5.6 mM KCl and 1.2 mM MgSO_4_, 1.2 mM KH_2_PO_4_, 11.1 mM glucose, 10 mM TES and 2.4 mM CaCl_2_). Each strip was set to a resting tension of 2g and contractile activity was recorded using isometric force transducers (DMT; Muscle Strip System −820MS). Data acquisition and analysis were performed using Lab Chart software (AD Instruments). Myometrium was left to equilibrate for up to 120 minutes until regular spontaneous contractions were observed. Experiments were performed only when four strips had regular contractions. Baseline contraction was recorded for 10 minutes, and myometrium was then either left to spontanesouly contract or stimulated with 10^−7^M oxytocin for 30 minutes. Myometrial strips were randomized and received two sequential doses, low (6 μM) and high (9 μM), of either UK5099 or etomoxir. Contractile activity was measured for 10 minutes after drug stimulation, including activity integrals (area under the force-time curve), mean amplitude and contraction frequency (peaks per minute) Viability of myometrial strips was confirmed at the end of each experiment by recording contractile responses to administration of 50 mM KCl.

### Flow cytometry

Flow cytometry was performed as previously described^59^, with minor modifications. Briefly, cells were transferred from 6-well plates to a round-bottom 96-well plate (ThermoFisher) by washing with Dulbecco’s Phosphate Buffered Saline (DPBS; Gibco) and lifting cells with TrypLE Express (Gibco). Next, 10% FBS was added to the wells before plating an equal amount of the cell suspension to each well with technical replicates. Cells were pelleted by centrifugation at 1000 RCF for 5 mins at 4 °C, washed with Fluorescent Activated Cell Sorting (FACS) buffer (DPBS, 0.5% BSA, 0.1% NaN_3_, pH 7.4), and stained on ice for 40 mins in FACS buffer with 0.1% Invitrogen eBioscience Fixable Viability Dye eFluor 780 (ThermoFisher). Cells were then washed twice in FACS buffer, resuspended in 100 μL fresh FACS buffer and analyzed at 4 °C on a Cytek Aurora Flow Cytometer (Cytek Biosciences). Samples were gated for cell size with forward scatter and side scatter height (FSC-H, SSC-H), then gated for single cells by side scatter height and forward scatter area (FSC-A, SSC-H), and then gated by FSC-A and the Viability cell stain to identify live versus dead cells. Cell viability was not affected by extended oxytocin treatment (Supplementary Figure S1).

### Statistical analysis

Experiments for mitochondrial and glycolytic function analysis were replicated in three or four wells. Results were averaged for each treatment group. Data are presented as means ± standard errors of the means (SEM). Significance was determined by repeated measures one-way ANOVA or two-way ANOVA, as appropriate. Holm-Sidak test was used to adjust for multiple comparisons. Prism 6 software (GraphPad) was used for statistical analyses. Significant differences compared to controls are annotated as follows: *P < 0.05, **P < 0.01, ***P < 0.001, and ****P < 0.0001.

### Study approval

Washington University in St. Louis Human Research Protection Office approved this study (IRB #202511061). Deidentified human myometrial samples were obtained under a waiver of consent from the Reproductive Specimen Processing and Banking core (ReProBank) at Washignton University in St. Louis.

## Data availability

The authors agree to share all publication-related data.

## AUTHOR CONTRIBUTIONS

AIF and KKP conceptualization; AIF funding acquisition; KKP, RMD, KTM, JLR, RMG, MNM and AIF methodology; KKP, RMD, KTM, JLR and AIF investigation and experimental work; KKP, RMD, KTM and AIF interpretation of results; KKP and AIF writing—original draft; KKP, RMD, KTM, JLR, RMG, MNM and AIF writing—review and editing. All authors read and approved the final version.

## ACKNOWLEDGEMENTS

We thank Dr. Deborah Frank for invaluable editorial assistance. We thank the Reproductive Specimen Processing and Banking core (ReProBank) for assistance with human specimen procurement. We thank the patients at Barnes Jewish Hospital for being part of this research study. Seahorse was performed and supported by the Washington University Diabetes Research Center (P30DK020579). This was was supported by NIH grant R21 HD110610 (A.I.F.).

