## Supplemental Data for "Metabolic Flexibility and Energy Substrate Utilization Regulate Contractility in the Human Myometrium"

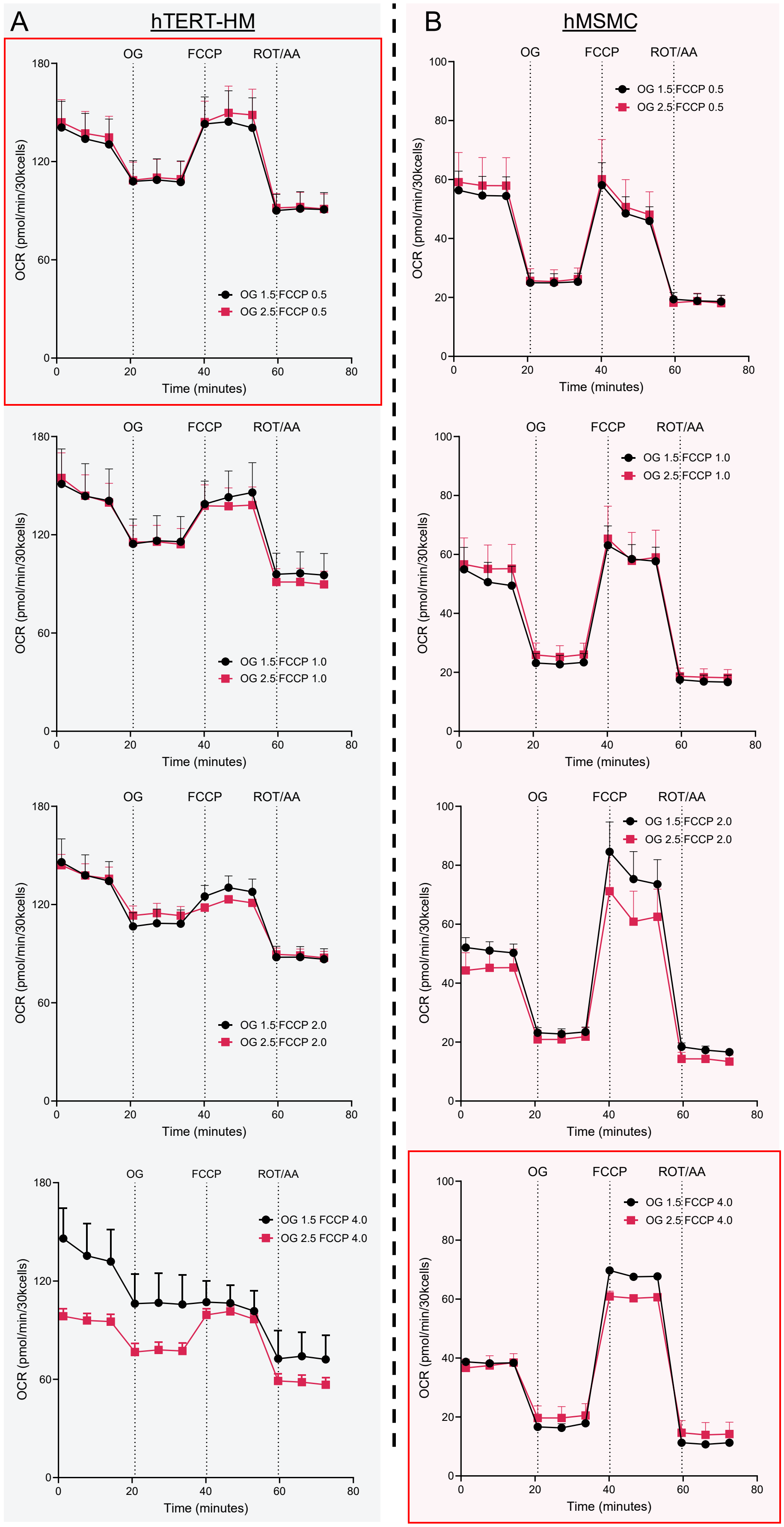


Supplementary Figure S1

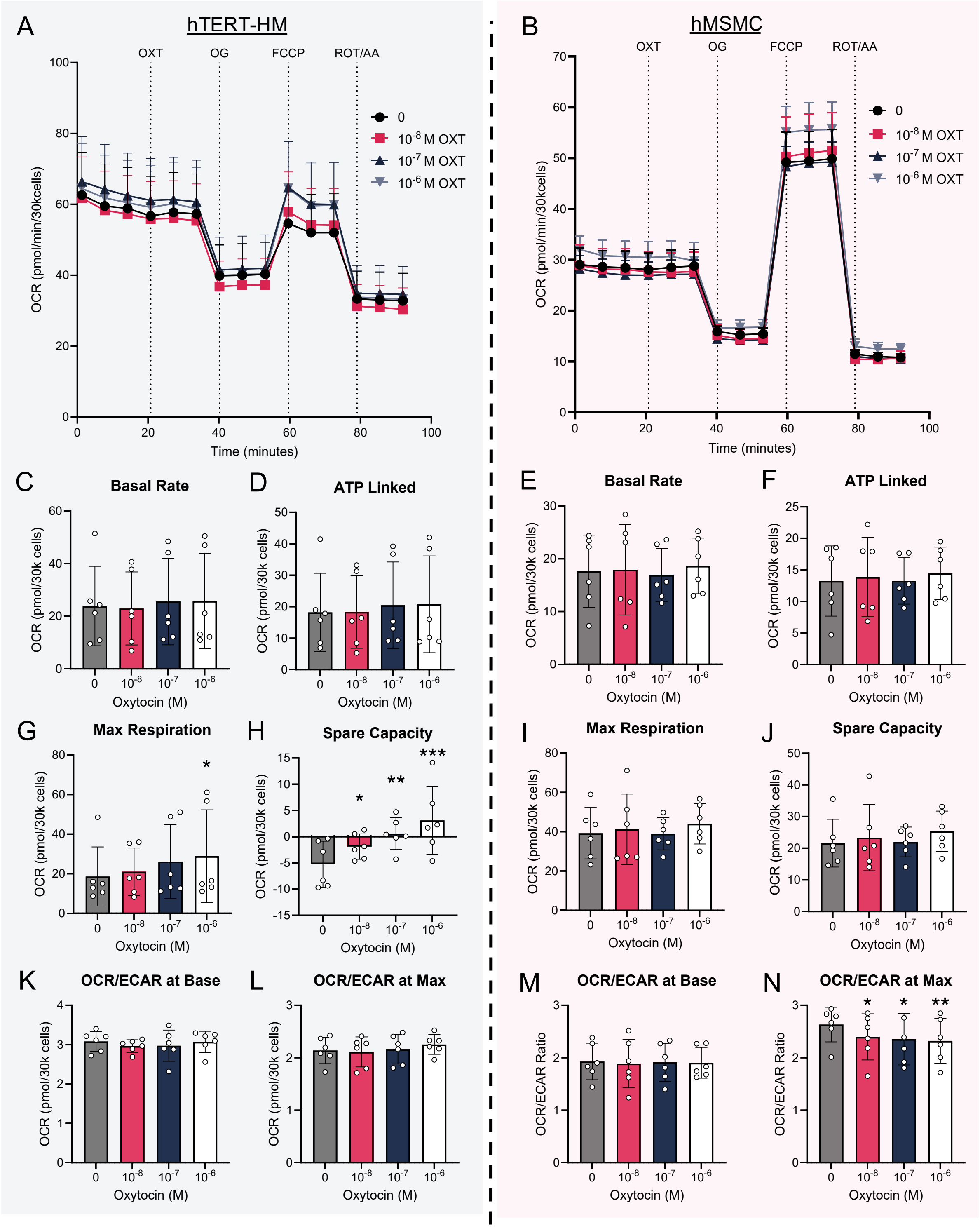


Supplementary Figure S2

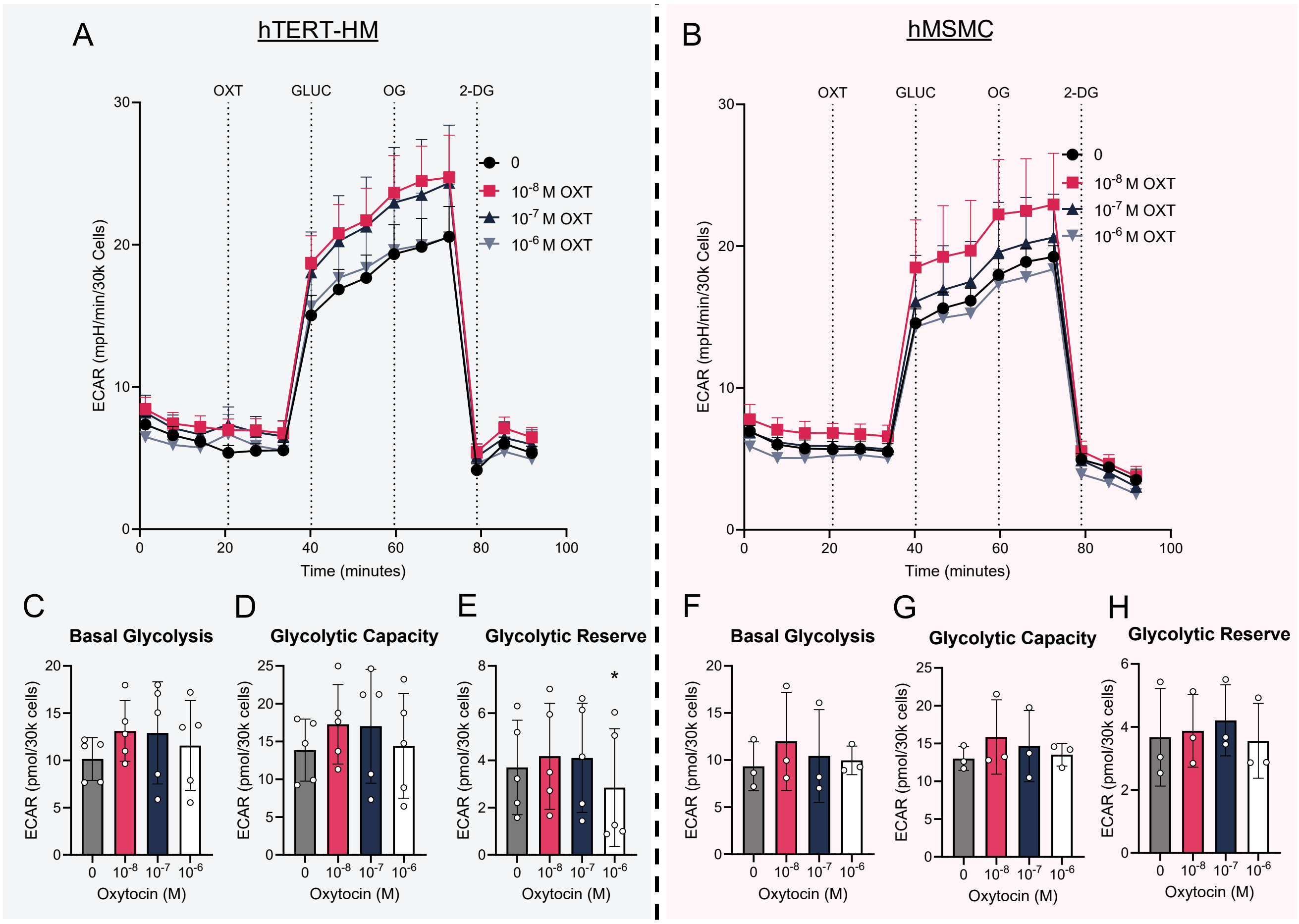


Supplementary Figure S3

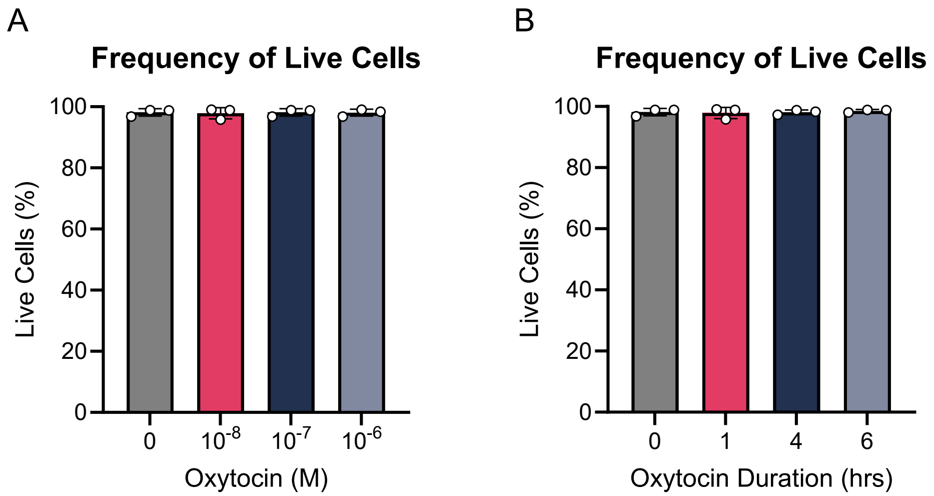


Supplementary Figure S4
